# Aberrant splicing prediction across human tissues

**DOI:** 10.1101/2022.06.13.495326

**Authors:** Muhammed H. Çelik, Nils Wagner, Florian R. Hölzlwimmer, Vicente A. Yépez, Christian Mertes, Holger Prokisch, Julien Gagneur

## Abstract

Aberrant splicing is a major cause of genetic disorders but its direct detection in transcriptomes is limited to clinically accessible tissues such as skin or body fluids. While DNA-based machine learning models allow prioritizing rare variants for affecting splicing, their performance on predicting tissue-specific aberrant splicing remains unassessed. Here, we generated the first aberrant splicing benchmark dataset, spanning over 8.8 million rare variants in 49 human tissues. At 20% recall, state-of-the-art DNA-based models cap at 10% precision. By mapping and quantifying tissue-specific splice site usage transcriptome-wide and modeling isoform competition, we increased precision by three-fold at the same recall. Integrating RNA-sequencing data of clinically accessible tissues brought precision to 60%. These results, replicated in two independent cohorts, substantially contribute to non-coding loss-of-function variant identification and to genetic diagnostics design and analytics.

## Main

Identifying non-coding loss-of-function variants is a major bottleneck of whole genome interpretation^1^. Variants altering splicing represent a large class of non-coding loss-of-function variants because they can lead to drastically altered RNA isoforms, for instance by inducing frameshifts or ablations of functionally important protein domains. If the variant effect is strong, the proportion of non-functional RNA isoforms can become so large that the function of the gene is lost. Due to the relevance of splicing for variant interpretation, notably in rare disease diagnostics and in oncology, algorithms have been developed to predict whether variants affect splicing^2–6^. However, only recently, aberrant splicing events, defined as rare and quantitatively large alterations of splice isoform usage, have been called in human tissues^7,8^. While a method to a posteriori prioritize candidate causal rare variants for observed aberrant splicing events has been proposed^7^, the forward problem, i.e. predicting among rare variants which ones cause aberrant splicing, has not been addressed.

Here, we set out to establish models predicting whether a rare variant associates with aberrant splicing in any given human tissue. First, we assumed only DNA to be available and later on further considered complementary RNA-sequencing (RNA-seq) data of clinically accessible tissues (Fig. 1).

**Fig. 1:**
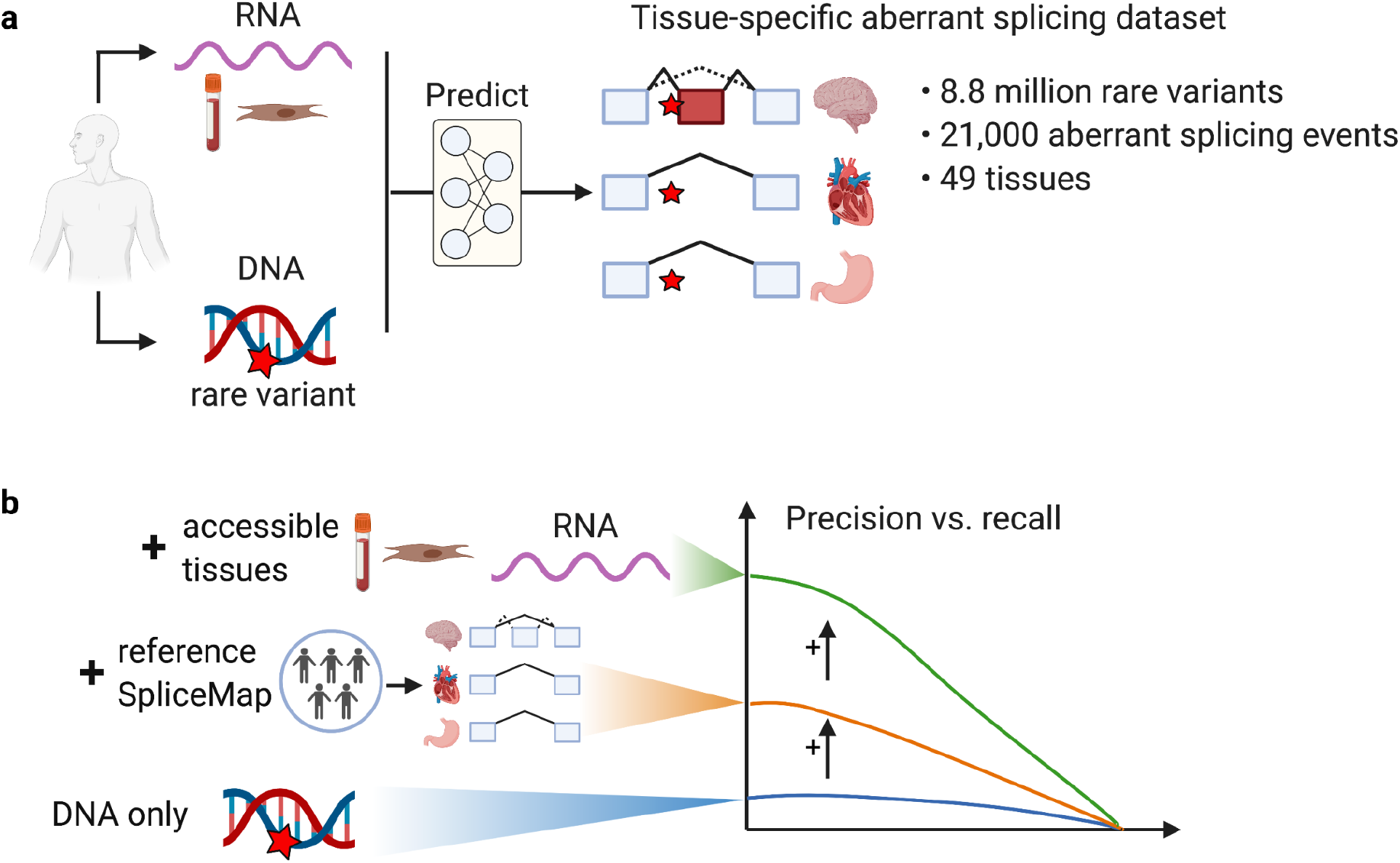
Study design and main findings. We set out to predict whether rare variants associate with aberrant splicing across 49 human tissues. **a**, We established the first benchmark for aberrant splicing by processing GTEx samples with a recently published aberrant splicing caller^8^ based on which we could assess and develop predictors that could take as input DNA sequence and, optionally, RNA sequencing data of clinically accessible tissues. **b**, Benchmarking revealed modest performance of currently used algorithms based on DNA only, a substantial performance improvement when integrating these models with SpliceMap, a quantitative map of tissue-specific splicing we developed in this study, and further improvements when also including direct measures of aberrant splicing in accessible tissues.

## Results

We created a benchmark using the aberrant splicing caller FRASER^8^ on 16,213 RNA-seq samples of the GTEx dataset, spanning 49 tissues and 946 individuals. For every individual, we considered every protein coding gene carrying at least one rare variant (minor allele frequency less than 0.1% based on gnomAD^9^ and found in no more than two individuals across GTEx) and set out to predict in which tissue, if any, is this gene aberrantly spliced. We defined a gene to be aberrantly spliced in a sample if it was called as a transcriptome-wide significant splicing outlier and with a sufficient amplitude (differential percent spliced-in larger than 0.3, Methods). Previous studies had reported that as many as 75% of aberrant splicing events in GTEx RNA-seq samples are not replicated across tissues^7,8^ and thus may reflect technical artifacts or aberrant splicing that is not genetically driven. Therefore, we also required a rare variant to be less than 100 nucleotides (nt) away from the boundaries of any intron associated with the aberrantly spliced splice site (Methods). This filter yielded similar results as filtering for replicated aberrant events with the extra advantage of being applicable to independent cohorts that have a single sample per individual (Supplementary Figs. 1-2).

We first assessed the performance of two complementary state-of-the-art sequence-based deep learning models: MMSplice^3^, which predicts quantitative usage changes of predefined splice sites, and SpliceAI^2^, which predicts creation or loss of splice sites anywhere in a gene. For individuals with multiple rare variants on a gene, we retained the largest score of each model. Out-of-the box application of MMSplice and SpliceAI showed a modest performance with an overall precision of 8% for MMSplice and of 12% for SpliceAI at 20% recall, and an area under the precision-recall curve (auPRC) of 4% ± 1 percentage points across tissues for MMSplice and 5% ± 2 percentage point for SpliceAI.

We observed that many false predictions originated from inaccurate genome annotations. On the one hand, standard genome annotations are not tissue-specific, leading to false positive predictions. This includes predictions for genes that are not expressed in the tissue of interest, as for the gene *TRPC6* in the brain (Fig. 2a) and, among expressed genes, predictions for exons that are not canonically used in the tissue, as for exon 2 of *C2orf74* in the tibial nerve (Fig. 2b). On the other hand, many splice sites are missing from standard genome annotations^10,11^. These non-annotated splice sites are often spliced at a low level, yet can be strongly enhanced by variants (see Fig. 2c for an example) and are suspected to be a major cause of aberrant splicing^12,13^. To address all these issues, we created a tissue-specific splice site map, which we named SpliceMap, using GTEx RNA-seq data. SpliceMap excludes untranscribed splice sites and introns for each tissue and includes non-annotated splice sites and introns reproducibly observed among samples of the same tissue (Methods). The standard genome annotation GENCODE^14^ contains 244,189 donor sites and 235,654 acceptor sites, of which 93% were detected at least in one GTEx tissue (Fig. 2d). SpliceMap contains 168,004 ± 9,288 donor sites and 164,702 ± 8,950 acceptor sites per tissue (Supplementary Fig. 3). From this total, 7,060 ± 3,706 donor sites and 8,222 ± 3,740 acceptor sites were novel, with testis containing the maximum number of non-annotated donor and acceptor sites (29,673 and 29,911 respectively), in line with the unique transcriptional and splicing pattern of testis^15,16^. Applying MMSplice on the tissue-specific splice sites defined by SpliceMap increased the precision of MMSplice to 11% at 20% recall (Fig. 2e) with a significantly higher auPRC consistently across tissues (Fig. 2f).

**Fig. 2:**
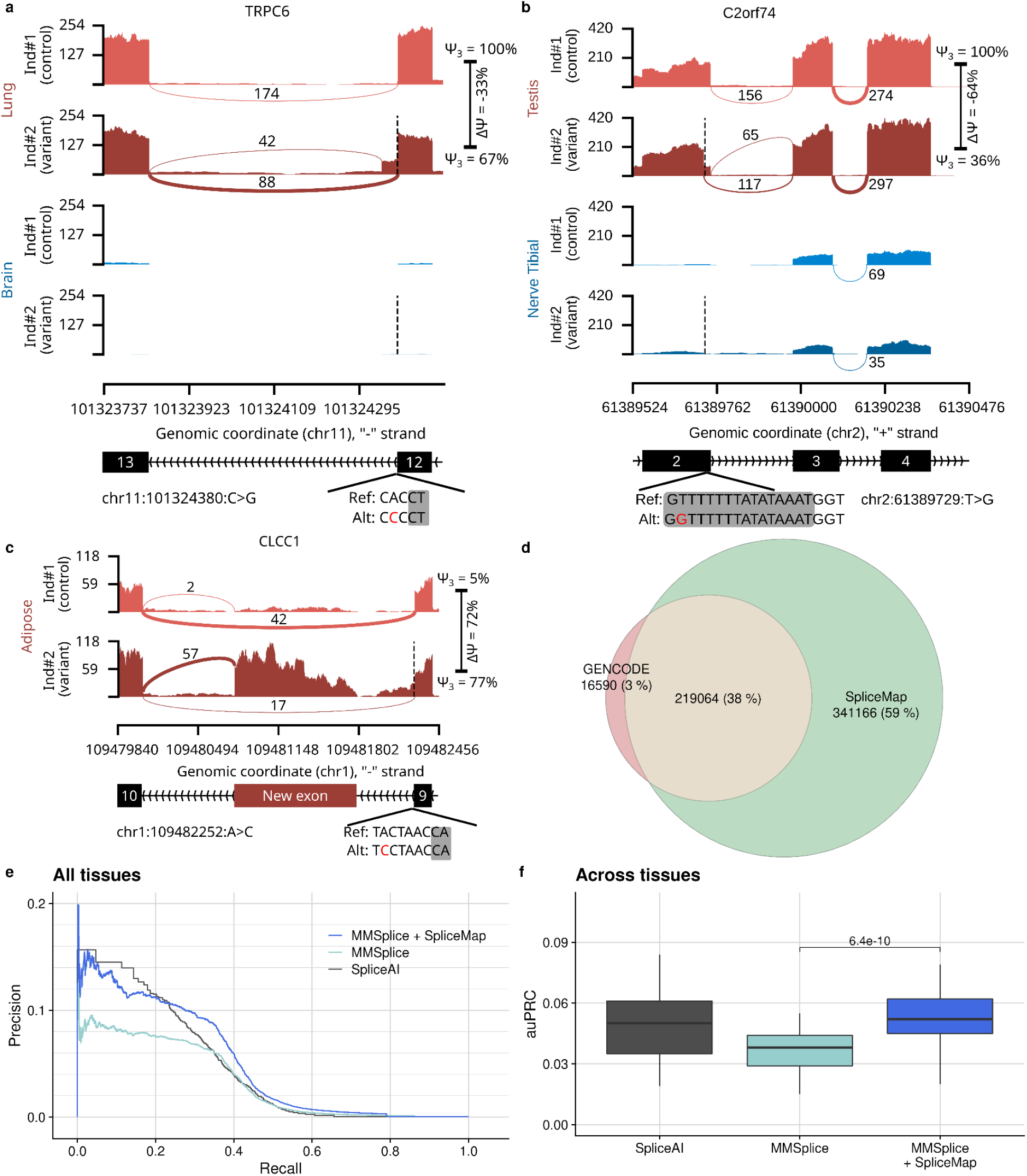
Tissue-specific splice site map improves prediction performance. **a-c**, So-called sashimi plots showing RNA-seq read coverage (y-axis) and the numbers of split reads spanning an intron indicated on the exon-connecting line (using pysashimi^17^) for instances illustrating the benefits of the SpliceMap annotation. For each instance, two individuals are displayed. The individual with the rare genetic variant (located at the dashed black line) is shown in the lower track (darker color). SpliceMap catalogs expressed genes and splice sites in each tissue and can thus help identifying cases for which there is no variant effect in tissues not expressing the whole gene **(a)** or the exon **(b)** in the proximity of the variant. Moreover, SpliceMap includes weak splice sites, which are spliced at a low level, but can be activated and create novel exons in the presence of a variant **(c). d**, Venn diagram comparing annotated splice sites in standard genome annotation (GENCODE release 38) and SpliceMap aggregating all GTEx tissues. **e**, Precision-recall curves comparing the overall prediction performance across all GTEx tissues (*n*=49) of MMSplice applied to GENCODE splice sites, MMSplice applied to tissue-specific splice sites according to SpliceMap, and SpliceAI. **f**, Distribution of the area under the precision-recall curve (auPRC) across all GTEx tissues of the models in (e). Center line = median; box limits = first and third quartiles; whiskers span all data within 1.5 interquartile ranges of the lower and upper quartiles. *P* values were computed using the paired one-sided Wilcoxon test.

Variants affecting splicing typically associate with abundance ratio fold-changes of competing splicing isoforms, which result in nonlinear effects on isoform proportions according to the so-called scaling law of splicing^18,19^. For instance, starting from a 1:1 ratio between one splicing isoform and its alternative in a major allele background, a ten-fold decrease leads to a 1:10 ratio, which amounts to around 40 percentage points decrease (from 50% to ca. 10%). However, the same ratio fold-change starting from a 1:10 ratio amounts to less than 1 percentage point decrease (Supplementary Fig. 4a). Hence, the scaling law of splicing implies that the sole variation of isoform abundance between tissues in major allele background can explain some of the tissue-specific effects of variants on isoform proportion^18^, as exemplified with the exon 7 of the gene *TRPC6* (Fig. 3a). We estimated major allele background levels of alternative donor and acceptor splice site usage proportions for all introns and all tissues of SpliceMap (Supplementary Fig. 4b-d). Integrating these reference levels further improved the MMSplice predictions by 1.6-fold consistently across tissues (Fig. 3b-c). Next, to leverage the complementarity of MMSplice and SpliceAI predictions^20^, we trained an integrative model using the scores from both deep learning models as well as annotation features from tissue-specific SpliceMaps (Methods). This model, that we call AbSplice-DNA, achieved an additional 1.5-fold improvement (Fig. 3b-c). The AbSplice-DNA scores are probability estimates which we found to be well-calibrated on GTEx (Supplementary Fig. 5). To ease downstream applications we suggest three cutoffs (high: 0.2, medium: 0.05, low: 0.01), which approximately have the same recalls as the high, medium and low cutoffs of SpliceAI (Fig. 3b).

**Fig. 3:**
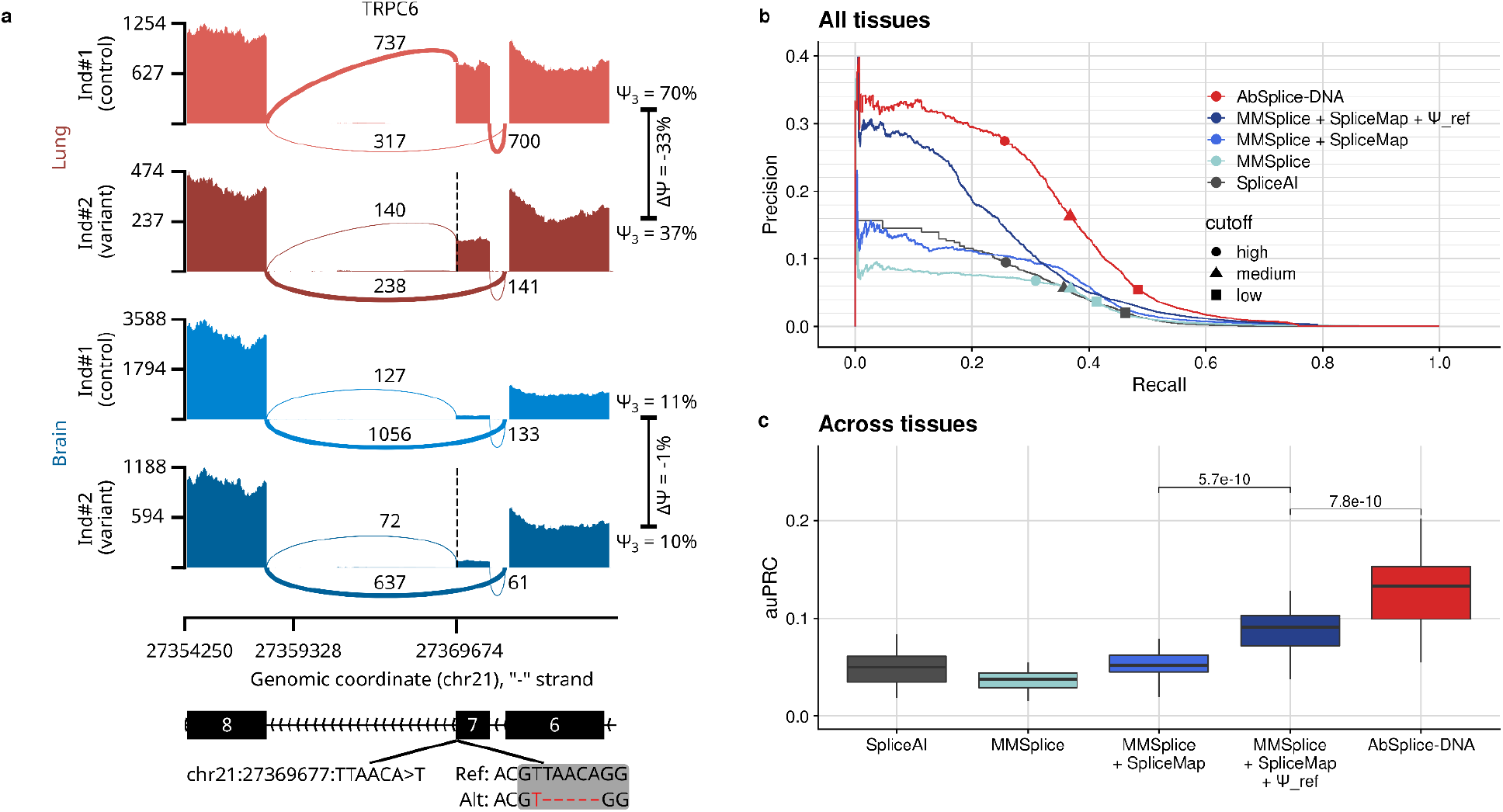
Quantitative splicing levels further improve prediction performance. **a**, Sashimi plot of *TRPC6* around exon 7 for two individuals in lung and brain, one carrying no rare variant in this region (control, upper tracks), and one carrying an exonic rare deletion (dashed line and lower tracks) associated with reduced splicing of exon 7. The donor sites of exon 6 and exon 7 compete against each other for splicing with the acceptor site of exon 8. For the control individual, the donor site of exon 7 is used 70% of the time in the lung, and only 11% of the time in the brain. The variant associates with a stronger difference (33 percentage points) in the lung than in the brain (1 percentage point). **b**, Precision-recall curve comparing the overall prediction performance on all GTEx tissues of SpliceAI, MMSplice using GENCODE annotation, MMSplice using SpliceMap annotation, MMSplice using SpliceMap annotation along with quantitative reference levels of splicing, and the integrative model AbSplice-DNA. Different cut-offs are shown (SpliceAI, high: 0.8, medium: 0.5, low: 0.2; MMSplice (score absolute value), high: 2, medium: 1.5, low: 1; AbSplice-DNA, high: 0.2, medium: 0.05, low: 0.01). **c**, Distribution of the area under the precision-recall curve of the models in (b) across tissues (*n*=49). Center line = median; box limits = first and third quartiles; whiskers span all data within 1.5 interquartile ranges of the lower and upper quartiles. *P* values were computed using the paired one-sided Wilcoxon test.

Having established our model on GTEx, we next assessed how the performance replicated in independent cohorts. We first evaluated a dataset consisting of RNA-seq samples from skin fibroblasts of 303 individuals with a suspected rare mitochondriopathy^21^. We found that there was a large overlap (86%) of splice sites in SpliceMaps generated from GTEx fibroblasts and from this cohort (Fig. 4a). Moreover, we observed highly consistent reference levels of splicing between the two datasets (Fig. 4b, R^2^=0.996). We applied AbSplice-DNA trained on GTEx using the SpliceMap from GTEx fibroblasts on the subset of this data for which whole genome sequencing was available (*n*=20) and used aberrant splicing calls performed on the RNA-seq samples to assess the predictions. The relative improvements between the baseline models and AbSplice-DNA replicated. AbSplice-DNA achieved 12.8% ± 1.3% auPRC, three-fold higher than SpliceAI or MMSplice alone (Fig. 4c). From a rare variant prioritization stand point, AbSplice-DNA typically gave about 2-fold less candidate predictions at the same level of recall than SpliceAI, itself comparing favorably over MMSplice (Fig. 4d). Hence, AbSplice-DNA can help rare disease diagnostics by providing substantially shorter lists of predicted candidate variants to investigate compared to state-of-the–art sequence-based models.

**Fig. 4:**
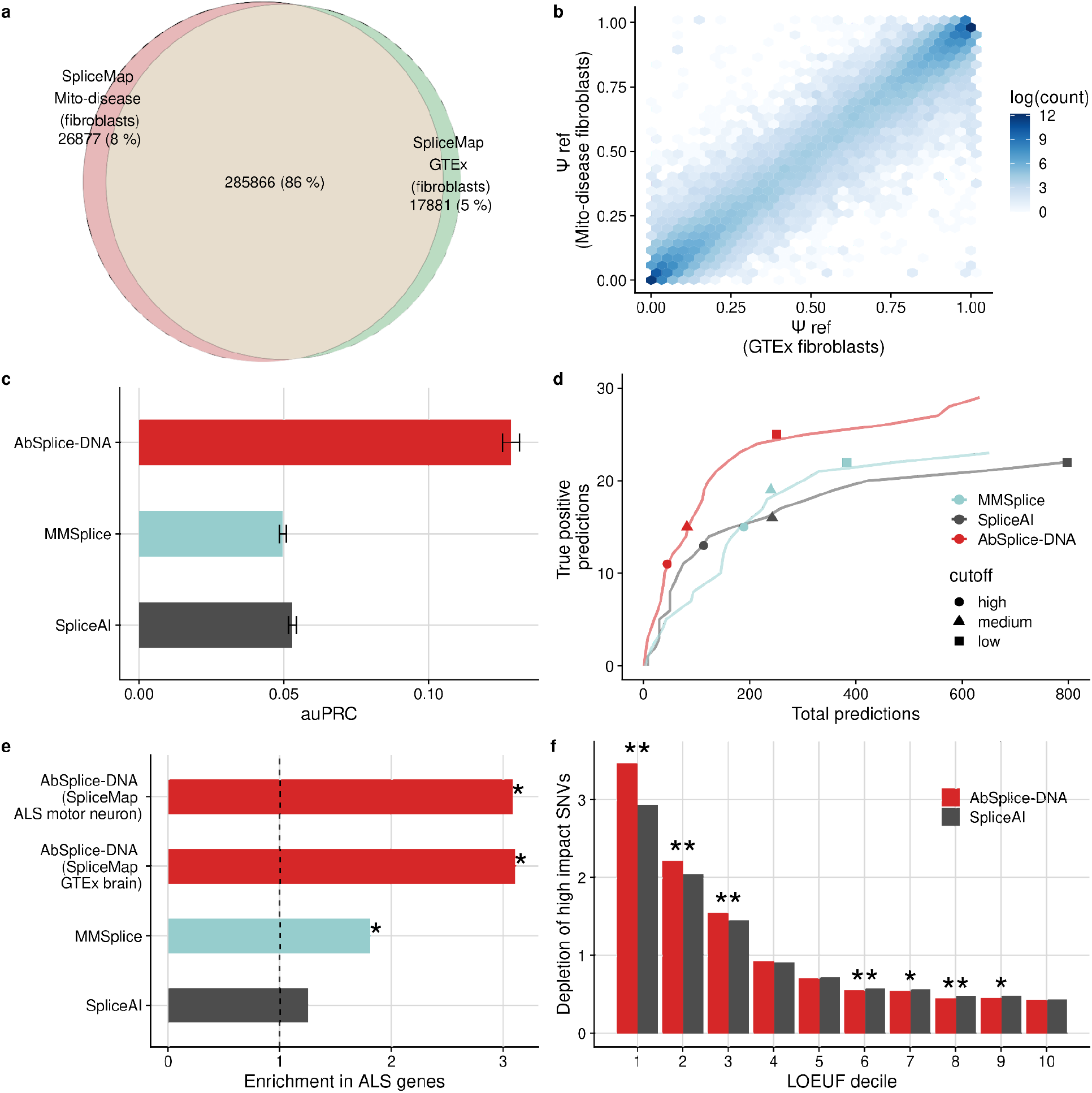
Application of AbSplice-DNA on independent data. **a**, Venn diagram comparing the splice sites in the SpliceMap generated from fibroblasts from a mitochondrial disease dataset (*n*=303) and GTEx (*n*=492). **b**, Relation of the reference Ψ values from the SpliceMaps from (a). **c**, AuPRC for classification of aberrant splicing events from rare variants in the mitochondrial disease dataset for SpliceAI, MMSplice and AbSplice-DNA trained on GTEx using the GTEx fibroblasts SpliceMap from (a). Error bars represent standard errors of the mean (Jackknife over samples). **d**, Variants predicted to cause aberrant splicing among splicing outliers against all predicted variants for different methods and showing different cut-offs (SpliceAI, high: 0.8, medium: 0.5, low: 0.2; MMSplice (absolute of score), high: 2, medium: 1.5, low: 1; AbSplice-DNA, high: 0.2, medium: 0.05, low: 0.01). **e**, Enrichment of high score predictions in ALS genes (*n*=165). Stars mark significant one-sided Fisher tests considering all protein coding genes as the universe (*P* < 0.05). **f**, Genome-wide depletion of high impact variants among rare SNVs (gnomAD MAF < 0.1%) within a gene (*n*=19,521) as a function of LOEUF score deciles. High impact variants are defined by a SpliceAI score > 0.8 and an AbSplice-DNA score > 0.2 in at least one tissue. Stars mark significance levels of two-sided Fisher tests (* < 0.05, ** < 10^−4^).

**Fig. 5:**
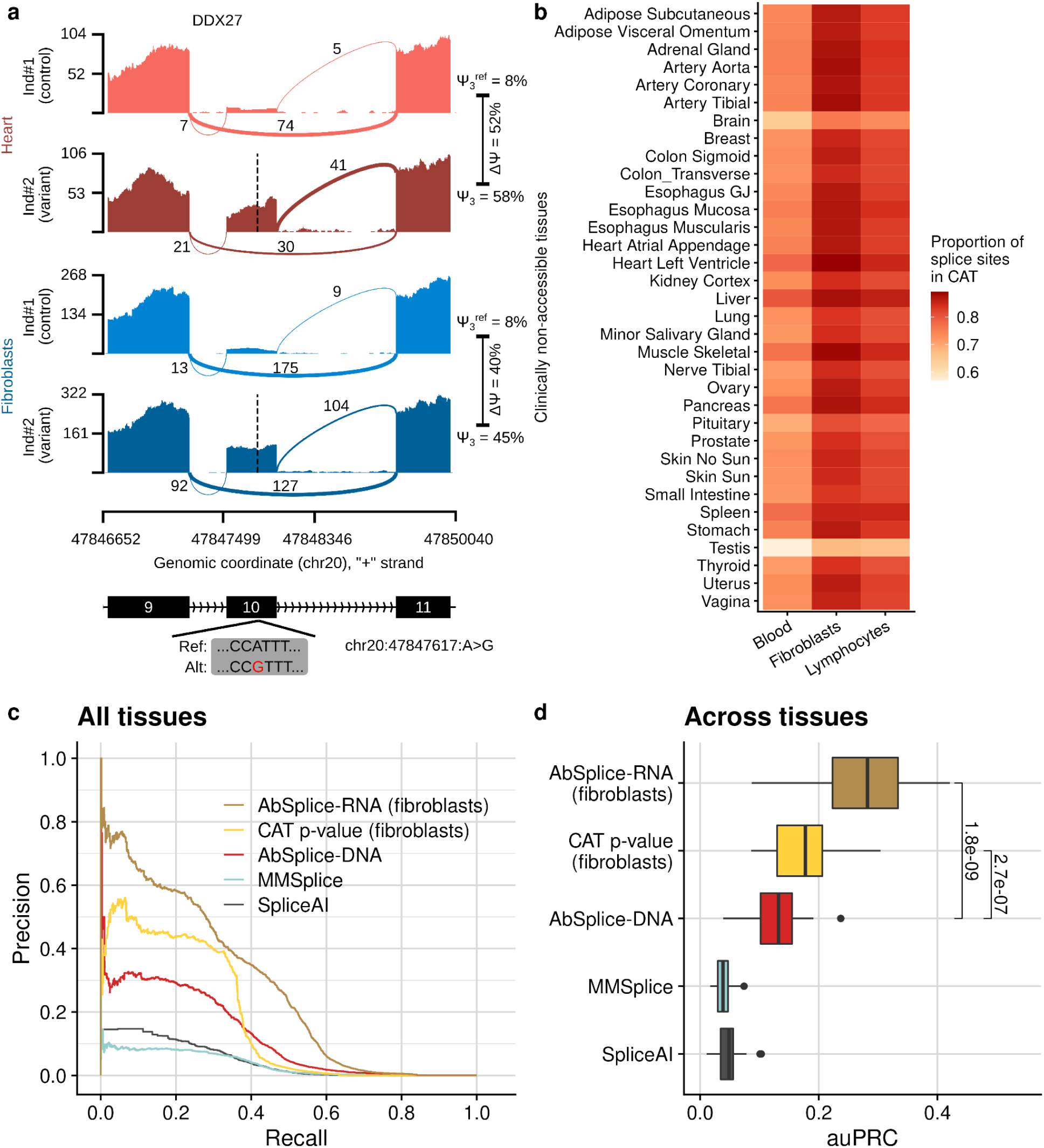
Integrating RNA-seq data of clinically accessible tissues to predict aberrant splicing in non-accessible tissues. **a**, Sashimi plot of *DDX27* around exon 10 for two individuals in heart and fibroblasts. One individual carries no rare variant in this region (control, upper tracks), and one carries an exonic rare variant (dashed line, lower tracks) associated with increased splicing of exon 10. This exon shows a similar usage in fibroblasts and in the heart (reference donor site percent spliced-in Ψ_5_ = 8% according to SpliceMap in both tissues, in line with the measured values for the displayed control individual). The effect associated with the variant in fibroblasts approximates well the one in heart (difference of donor site usage ΔΨ_5_ = 50% in heart and 37% in fibroblasts). In this case, aberrant splicing can be directly detected from the accessible tissue. **b**, Proportion of splice sites used in clinically non-accessible target tissues (rows) also used in CATs (columns). **c**, Precision-recall curve comparing the overall prediction performance on all GTEx tissues of SpliceAI, MMSplice using GENCODE annotation, AbSplice-DNA, gene-level FRASER p-values in fibroblasts, and AbSplice-RNA which integrate AbSplice-DNA features with features from RNA-seq from fibroblasts. **d**, Distribution of the area under the precision-recall curve of the models in c) across tissues (*n*=49). Center line = median; box limits = first and third quartiles; whiskers span all data within 1.5 interquartile ranges of the lower and upper quartiles; *P* values were computed using the paired one-sided Wilcoxon test.

We next considered a cohort of whole genome sequencing samples paired with RNA-seq data of iPSC-derived spinal motor neurons from 245 Amyotrophic lateral sclerosis (ALS) affected and 45 healthy individuals from the Answer ALS project (Methods). As motor neurons were not profiled in GTEx, we considered two approaches. On the one hand, we used the Answer ALS healthy controls to generate a SpliceMap for motor neurons. On the other hand, we used the SpliceMap of GTEx brain tissues as a proxy. We found that the GTEx SpliceMap from brain tissues agreed reasonably well with the one derived from this cohort both qualitatively (76% shared splice sites) and quantitatively (R^2^=0.98, Supplementary Fig. 6a-b). Here too, AbSplice-DNA outperformed SpliceAI and MMSplice. Interestingly AbSplice-DNA achieved similar performances using the SpliceMap from GTEx brain tissues or using the SpliceMap from motor neurons, suggesting that AbSplice-DNA can be applied robustly in absence of control samples using SpliceMaps from proxy tissues (Supplementary Fig. 6). Moreover, AbSplice-DNA predictions were enriched for genes associated with ALS, which was less so for MMSplice predictions and not the case for Splice-AI predictions (3-fold enrichment, Fig. 4e).

Furthermore, we applied AbSplice-DNA to 199,489,923 rare variants (MAF < 0.1%) from the gnomAD datasets. In highly constrained genes, defined as the 10% genes most strongly depleted for loss-of-function variants in gnomAD^9^, rare variants were more strongly depleted for high AbSplice-DNA scores in at least one tissue (3.5-fold depletion), than for high SpliceAI scores (2.9-fold depletion, *P*<10^−21^, Fig. 4f). A stronger depletion than with SpliceAI also held when relaxing the AbSplice-DNA cutoff to match the total number for predictions of SpliceAI (Supplementary Fig. 7).

Collectively, these results on independent data demonstrate the robustness and the applicability of AbSplice-DNA and indicate its relevance for rare disease diagnostics and rare variant interpretation.

Sequencing transcriptomes of clinically accessible tissues (CATs) such as skin or body fluids is of increasing interest in rare disease research as it allows direct detection of aberrant splicing for those splice sites used both in the CAT and in tissues of suspected disease relevance^12,22–24^. The GTEx dataset consists of post-mortem collected RNA-seq samples across a vast variety of tissues and thereby offers the unique opportunity to evaluate to what extent aberrant splicing in an accessible tissue reflects aberrant splicing of another tissue of interest. One positive example in GTEx is aberrant splicing of *DDX27* in the heart which can also be observed in skin fibroblasts (Fig. 5a). Consistent with a previous study^24^ based on the Ensembl gene annotation^25^, we found that among the CATs, fibroblasts have the highest overlap of transcribed splice sites according to SpliceMap with non-accessible tissues, followed by lymphocytes and whole blood (Fig. 5b). To predict aberrant splicing in non-accessible tissues, we first considered ranking genes of an individual first for showing significant and large aberrant splicing in a CAT (FDR <0.1 and Delta-Psi > 0.3) and then by significance level. This simple method yielded a markedly increased precision compared to the DNA-based models up to nearly 40% recall (Fig. 5c, Supplementary Fig. 8a). However, RNA-based predictions remain limited to those splice sites expressed and spliced in the CAT. Therefore, we next trained models integrating AbSplice-DNA features together with RNA-seq based features from CATs, including differential splicing amplitude estimates to leverage the splicing scaling law and the SpliceMaps (Methods). These models, that we call AbSplice-RNA, outperformed all other models (Fig. 5c and Supplementary Fig. 8a). We found that using fibroblasts only led to the same performance as using all CATs reaching around 60% precision at 20% recall and amounting to a 2-fold improvement over AbSplice-DNA (Fig. 5c and Supplementary Fig. 8b). Those improvements were consistent across target tissues (Fig. 5d). These results establish for the first time a formal way to integrate direct measurements of aberrant splicing along with sequence-based models to predict aberrant splicing in a tissue of interest.

## Discussion

We established the first benchmark for predicting variants leading to aberrant splicing in human tissues, revealing limited performance of state-of-the-art sequence-based models. We created a tissue-specific splicing annotation (SpliceMap) based on GTEx which maps acceptor and donor splices and quantifies their usage in 49 human tissues. We showed that integrating SpliceMap with DNA-based prediction models leads to a three-fold increase of precision at the same recall. Additionally, we found that RNA-seq from clinically accessible tissues complements DNA-based splicing predictions when incorporated into an integrative model, the first of this kind.

As large-scale cohorts of whole genome sequencing are on the rise in research and healthcare, there is an increasing need for efficient annotations of non-coding variants. Among those, variants that have strong deleterious effects are of special interest as they can contribute to effector gene identification and improved polygenic risk scores spanning the full spectrum of variant frequency. In this context, our model AbSplice-DNA not only outperforms state-of-the-art models, but it also provides tissue-specific predictions. Using aberrant splicing predictions for tissues that are mechanistically related to the disease of interest may prove to be helpful, just as tissue-specific predictions are important for transcriptome-wide association studies^26^.

We showed how RNA-seq of clinically accessible tissues effectively complement DNA-based predictions. An alternative to this approach is to reprogram or transdifferentiate cells into the suspected mechanistically-involved cell type and perform RNA-seq on them^27^. This approach has however important caveats. First, it is not ensured that the suspected mechanistically-involved cell type is the correct one as symptoms may manifest most strongly in downstream affected tissues. Second, this approach is labor-intensive. Third, cell reprogramming can induce and select mutations which may lead to false identifications. Therefore, predictive models that can leverage RNA-seq of CATs will probably remain relevant in practice.

By increasing the precision at 20% recall from about 10% to 60%, the cumulated improvements of our models are substantial. Still, a majority of the aberrant splicing events are not recalled and there remains a majority of false positives. An unknown and potentially large fraction of events that are not recalled might be aberrant splicing calling artifacts, as suggested by the high number of singleton calls. Progress in aberrant splicing calling or better understanding of the technical reasons could reduce the number of aberrant splicing events and improve the recall. Moreover, some of the apparent false positive predictions may be actually correct. This is the case when the aberrant splicing isoform contains a premature termination codon and, therefore, gets rapidly degraded by nonsense-mediated decay (NMD). Rapidly degraded isoforms barely lead any reads in RNA-sequencing data and hence are typically not detected by aberrant splicing callers. In diagnostic applications, those variants remain relevant. Moreover, dedicated experiments can be done to test whether aberrant splicing is taking place for instance using the translation inhibitor cycloheximide.

## Methods

### Datasets

#### GTEx

We downloaded the RNA-seq read alignment file (BAM files) and the variant calling files (VCF files) from whole genome sequencing (WGS) from GTEx v8p from dbGaP (study accession: phs000424.v8.p2). We used data from 946 individuals that have paired WGS and RNA-seq measurements (N=16,213) in at least one tissue.

#### Mitochondrial disease dataset

The dataset consists of 303 mitochondriopathy patients described in Yépez et al.^21^, all of which have RNA-seq from skin-derived fibroblasts. For 20 individuals, WGS is also available.

#### Amyotrophic Lateral Sclerosis (ALS) dataset

The dataset consists of WGS and RNA-seq from 245 individuals diagnosed with ALS and 45 control samples. RNA-seq data were obtained from iPSC derived spinal motor neurons. We downloaded the data from the Answer ALS portal (dataportal.answerals.org). Genes known to be involved in ALS disease development were manually curated from literature^28–31^.

### Data preprocessing

#### Rare variants

Variants had to be supported by at least 10 reads and had to pass the conservative genotype-quality filter of GQ ≥ 99. We considered a variant to be rare if it had a minor allele frequency (MAF) in the general population ≤ 0.001 based on the Genome Aggregation Database (gnomAD v3.1.2) and was found in at most 2 individuals within each cohort.

#### Splicing outlier detection

Splicing outliers were called using FRASER^8^ as implemented in the Detection of RNA-seq Outliers Pipeline^32^. FRASER was used to detect introns (including *de novo* introns) and to count split reads for each intron. Based on the split read counts, two intron-centric metrics were calculated: alternative acceptor usage with the Ψ_5_ alternative donor usage with the Ψ _3_ metric:

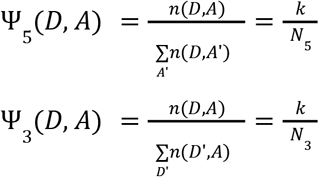

where *k* is the number of split reads supporting the intron from donor *D* to acceptor *A*. The sum in the denominator of Ψ_5_ (*D, A*) goes over all possible acceptors *A*’ for donor *D*, and the sum in the denominator of Ψ_3_ (*D, A*) goes over all possible donors *D*’ for acceptor *A*. The advantage of these intron-centric metrics over the exon-centric metric percent spliced-in (Ψ) is that they do not require exons to be mapped, which is an ill-defined task when starting from short-read RNA-sequencing data.

FRASER models these metrics while controlling for latent confounders and reports splice-site and gene-level splicing outliers with a multiple testing corrected p-value (Benjamini–Yekutieli FDR^33^). On the gene-level we selected outliers with a FDR-corrected p-value smaller than 0.1. We furthermore filtered for genes that also had a significant splice-site level outlier with FWER-corrected p-value smaller than 0.05, supported by 20 reads and with an absolute deviation of Ψ_5,3_ from the FRASER-modeled expected value larger than 0.3 (denoted |ΔΨ_5,3_| > 0. 3).

To discard aberrant splicing calls that have probably no genetic basis^8^, we additionally applied and compared different filtering methods (Supplementary Fig. 2). In the GTEx dataset, where multiple RNA-seq samples from the same individual are available, we investigated including splicing outliers found in RNA-seq data of at least two tissues from the same individual (filter 2, Supplementary Fig. 2). As this strategy could not be applied to other datasets, we alternatively filtered for splicing outliers containing a rare variant in the vicinity of ± 100 *bp* of the splice sites based on RNA-seq from the sample (filter 3). For consistency, all reported results are based on filter 3.

### Aberrant splicing prediction benchmark

#### Aberrant splicing prediction task

The task is to predict whether a protein-coding gene with one or more rare variants within the gene body is aberrantly spliced in a given tissue of an individual. *Performance evaluation metric*. Due to the large class imbalance in the splicing outlier prediction benchmarking dataset, we chose to evaluate models using precision-recall curves. As evaluation metric we used the area under the precision-recall curve, computed using the average precision score^34^ (which represents the mean of precisions for each threshold weighted by the recall difference):

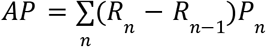

, where *P*_*n*_ and *R*_*n*_ are the precision and recall at the n^th^ threshold.

### Tissue-specific SpliceMap

For each tissue separately, we created a SpliceMap that lists all active introns along with aggregate statistics about acceptor and donor site usage useful for aberrant splicing prediction purposes.

#### Active introns

We started from all introns reported by FRASER. We filtered out untranscribed splice sites and background noise by filtering out introns not supported by any split read in more than 95% of the samples. For this and other operations involving genomic ranges we used PyRanges^35^.

#### Aggregate statistics

Aggregate statistics were calculated on donor and acceptor sites independently. For donor site usage, the SpliceMap aggregate statistics are i) the total number of split reads across samples (*s*) supporting the intron (Σ_*s*_*k*), ii) the total number of split reads across samples sharing the same acceptor site (Σ_*s*_*N*_3_), iii) the median number of split reads per sample sharing the same acceptor site, and iv) the reference isoform proportion (Ψ_3_^*ref*^) defined as 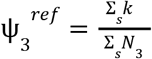. Aggregate statistics were computed analogously for acceptor site usage.

#### Exclusion of rare variant data in aggregate statistics

To prevent information leakage, the aggregate statistics were computed so that they do not contain information about splicing events associated with rare variants (specifically, we excluded from the computations of the aggregate statistics data from samples with a rare variant in ± 100 *bp* of any donor or acceptor site).

### Aberrant splicing prediction models

#### SpliceA

SpliceAI is a deep learning model that predicts splice site alteration for acceptor and donor sites from sequence^2^. SpliceAI is annotation free and can therefore score all variants including cryptic splice sites created by deep intronic variants. SpliceAI provides pre-computed scores for all SNVs and indels up to the length of four nucleotides. We downloaded pre-computed variant scores from Illumina BaseSpace and stored them in a RocksDB^36^ key-value database for fast look-up. We ran SpliceAI to obtain variant scores for long indels not available in the database. Also, we used masked scores of SpliceAI as recommended by the authors for variant interpretation. This masking sets Delta scores to zero if SpliceAI predicts strengthening for annotated splice sites and weakening for not-annotated splice sites.

#### MMSplice

MMsplice is a deep learning model that predicts the impact of a variant (in a 100 bp window of annotated splice sites) on alternative usage of a nearby donor or acceptor site^3^.

MMSplice predicts the effect of a variant in log-odds ratios (denoted Δ*logit* Ψ_5_ or Δ*logit* Ψ_3_).

MMSplice requires a splice site annotation. By default, we used the GENCODE (release 38) annotation.

#### MMSplice + SpliceMap

We ran MMSplice on tissue-specific splice site annotations from SpliceMap.

#### MMSplice + SpliceMap + Ψ_ref_

For conversion of the variant effect into natural scale, reference levels of donor site and acceptor site usages are required. For the sake of shorter notations, we write in the following Ψ instead of Ψ_5_ and Ψ_3_. We used MMSplice to predict Δ*logit* Ψ values. Δ*logit* Ψ values were then combined with the corresponding reference Ψ value (Ψ_*ref*_) in SpliceMap:

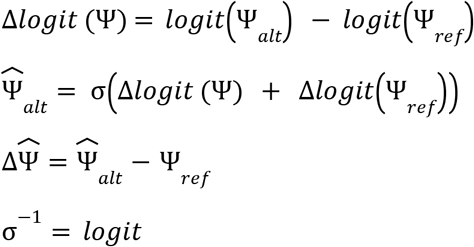

Variants further away than 100 nt from any SpliceMap splice site were scored 0 (no effect).

#### AbSplice-DNA

AbSplice-DNA is a generalized additive model, namely the ExplainableBoostingClassifier from the python package interpretml^37^. The features of AbSplice-DNA were the prediction score from MMSplice + SpliceMap, MMSplice + SpliceMap + Ψ_*ref*_, the SpliceAI Delta score, and a binary feature from SpliceMap indicating if the splice site is expressed in the target tissue (using a cutoff of 10 reads for the median number of split reads sharing the splice site). The model was trained with 5-fold stratified cross validation, grouped by individuals to avoid information leakage, and such that the proportion of the negative (no outlier on the gene) and positive (outlier on the gene) classes were preserved in each fold.

#### Predictors using RNA-seq from clinically accessible tissues (CAT)

We used different features from RNA-seq of three CATs from GTEx (Whole Blood, Cells transformed fibroblasts, and Cells Epstein-Barr virus (EBV) transformed lymphocytes) to predict aberrant splicing in non-accessible target tissues.

As one predictive feature we used the -log_10_ nominal gene-level p-values obtained using FRASER. In the benchmark, we ranked all splicing outlier genes (FDR < 0.1 and |ΔΨ | > 0. 3) lower than the remaining genes, and further ranked genes within each of these two groups by increasing p-value.

Additionally we used SpliceMaps from the accessible and the non-accessible tissues together with Ψ measurements from RNA-seq and applied the splicing scaling law to infer ΔΨ values in the non-accessible target tissue:

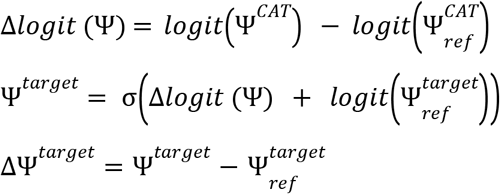

where = Ψ^*CAT*^ is the splicing level in the CAT, 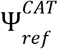 is the reference level of splicing obtained from SpliceMap and the difference of these two values provides the tissue unspecific variant effect Δ*logit* (Ψ). Then, adding of Δ*logit* (Ψ) with the reference level of splicing of the target tissue 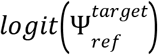 in logit scale and converting back to natural scale provides Ψ^*target*^ in the target tissue. Subtracting the reference level of splicing of the target tissue 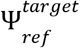 provides the predicted splicing change in the target tissue ΔΨ^*target*^ using RNA-seq measurements in CAT.

All precision-recall curves involving CATs have been computed on a subset of the data, excluding CATs from the target tissues and only containing individuals that have RNA-seq measurements from multiple tissues (including the CAT).

#### AbSplice-RNA

We trained integrative models using the two predictors from RNA-seq data from CATs described above in addition to DNA-based features used in AbSplice-DNA.

We trained AbSplice-RNA models using a single CAT and all CATs together. For the model using all CATs together we trained AbSplice-RNA in a CAT-agnostic manner such that the model predicts outliers regardless of the CAT source. This might be helpful in a diagnostic setting as it might be that the available CAT differs from the CATs that AbSplice-RNA was trained on.

#### Gene level aggregation

For genes with multiple variants, we retained the largest score per model.

### Enrichment in known ALS genes

The enrichment of 165 manually curated genes involved in ALS^28,29,31,38^ was computed as the proportion of high splicing impact variants within those genes, divided by all the high score predictions of the respective models. Depletion was computed as 1/enrichment.

### Depletion in loss-of-function intolerant genes

For all possible rare SNVs (gnomAD MAF < 0.1%) in 19,521 protein coding genes, we computed AbSplice-DNA scores and obtained the SpliceAI precomputed scores from Illumina BaseSpace. LOEUF scores were downloaded from https://gnomad.broadinstitute.org/downloads. For each LOEUF decile we computed the proportion of high splicing impact variants to the total sum of high impact variants and divided it by the proportion of rare variants in each decile.

## Code availability

SpliceMaps can be generated using the custom written python package ‘splicemap’ (publicly available at: https://github.com/gagneurlab/splicemap). AbSplice predictions using the enhanced SpliceMap annotation can be performed with the custom written python package ‘absplice’ (publicly available at: https://github.com/gagneurlab/absplice). We also provide a fast implementation of computing SpliceAI predictions using a wrapper based on fast lookup from a database of precomputed scores for existing variants and running SpliceAI for novel variants at https://github.com/gagneurlab/spliceai_rocksdb. Fast lookup of all gnomAD variants can be performed with https://github.com/gagneurlab/gnomad_rocksdb.

## Data availability

SpliceMaps for all 49 GTEx tissues and motor neurons from ALS (hg38) are available at Zenodo DOI: 10.5281/zenodo.6387937. Precomputed AbSplice-DNA scores (hg38) in all 49 GTEx tissues are available at Zenodo DOI: 10.5281/zenodo.6408331. Due to privacy restrictions we will approach the GTEx consortium to share the benchmark dataset upon acceptance of the manuscript.

## Acknowledgements

MHC thanks Xiaohui Xie and Ali Mortazavi for institutional support. The German Bundesministerium für Bildung und Forschung (BMBF) supported the study through the Model Exchange for Regulatory Genomics project (MERGE; 031L0174A to FRH and JG). NW is supported by the Helmholtz Association under the joint research school “Munich School for Data Science - MUDS”. This study was funded by the Deutsche Forschungsgemeinschaft (DFG, German Research Foundation) – via the projects “Identification of host genetic variation predisposing to severe COVID-19 by genetics, transcriptomics and functional analyses” (#466168909 to VAY and JG) and NFDI 1/1 “GHGA - German Human Genome-Phenome Archive” (#441914366 to CM and JG). Figure 1 was created with BioRender.com. The Genotype-Tissue Expression (GTEx) Project was supported by the Common Fund of the Office of the Director of the National Institutes of Health, and by NCI, NHGRI, NHLBI, NIDA, NIMH, and NINDS. This study was supported by data provided by the Answer ALS Consortium – administered by the Robert Packard Center for ALS at Johns Hopkins.

## Author contributions

Conceptualization: JG; Methodology: MHC, NW, JG; Software: MHC, NW; Validation: MHC, NW, FRH, VAY; Formal analysis: MHC, NW, FRH, VAY, CM; Data Curation: MHC, NW, FRH, VAY; Writing - Original Draft: MHC, NW, VAY, JG; Writing - Review & Editing: all authors; Visualization: MHC, NW, FRH, VAY, JG; Supervision: JG

## Competing interests

None

## Notes

https://zenodo.org/record/6631476

https://zenodo.org/record/6408906

https://github.com/gagneurlab/absplice

https://github.com/gagneurlab/splicemap

https://github.com/gagneurlab/spliceai_rocksdb

https://github.com/gagneurlab/gnomad_rocksdb

### Competing Interest Statement

The authors have declared no competing interest.

