## Supplementary Information for "Aberrant splicing prediction across human tissues"

### Aberrant splicing prediction across human tissues – Supplementary Information

#### Supplementary figures

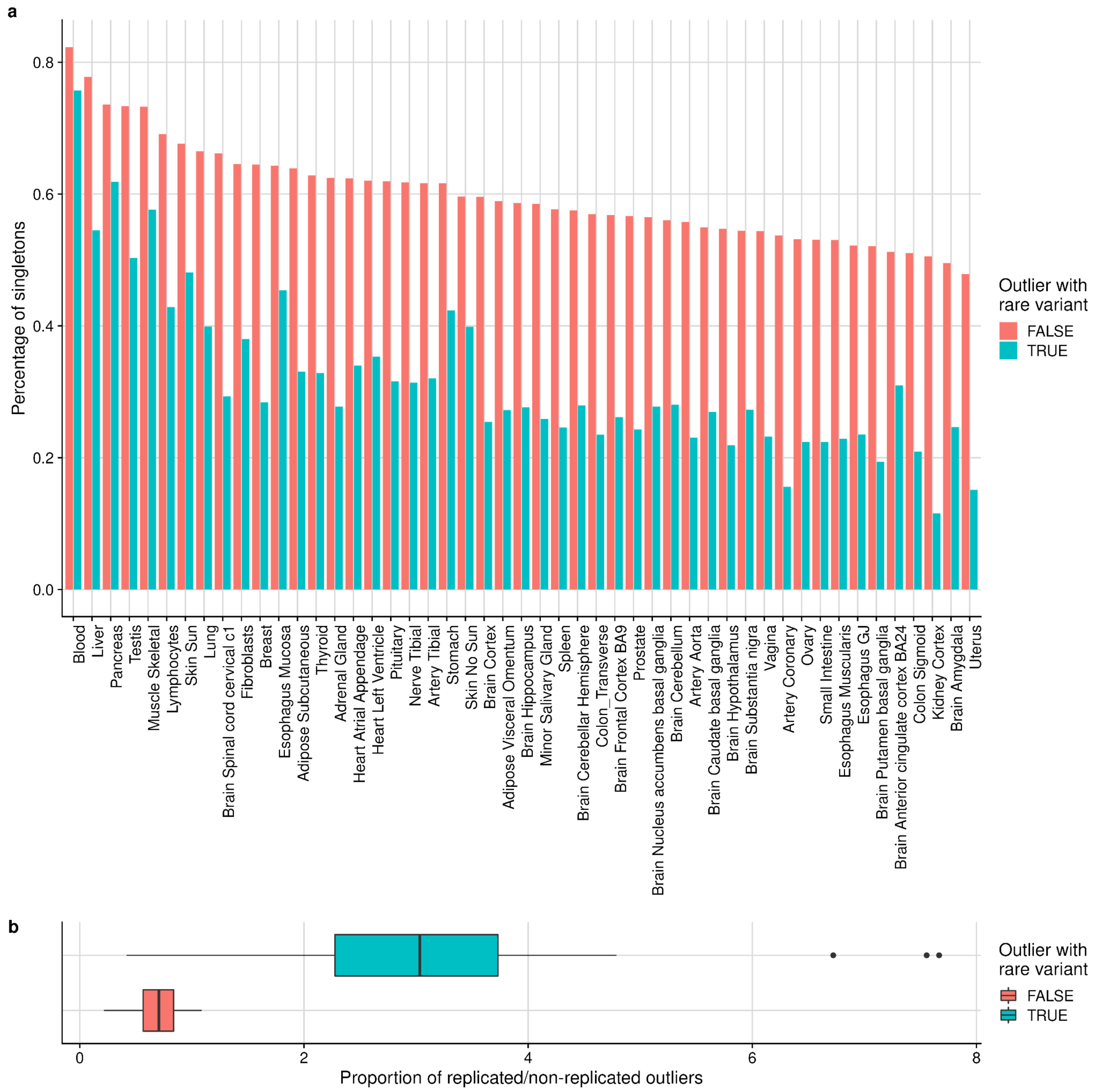


**Supplementary Fig. 1: Reduction of FRASER singletons by restricting to rare variants. a,** Percentage of singletons (aberrant splicing events that are observed only in one tissue) among all outliers (in red) and outliers with a rare variant (in blue) for each tissue. There are nearly no replicated RNA-seq samples in the GTEx dataset. Therefore, among all singleton events, genuinely tissue-specific aberrant splicing events are hard to distinguish from non-reproducible technical artifacts. **b**, Replication rate of aberrant splicing events between tissues of a sample for all aberrant splicing events (red) compared with aberrant splicing events that contain a rare variant in the vicinity (blue). Filtering for aberrant splicing events with a rare variant reduces the amount of singletons probably by filtering out technical artifacts.

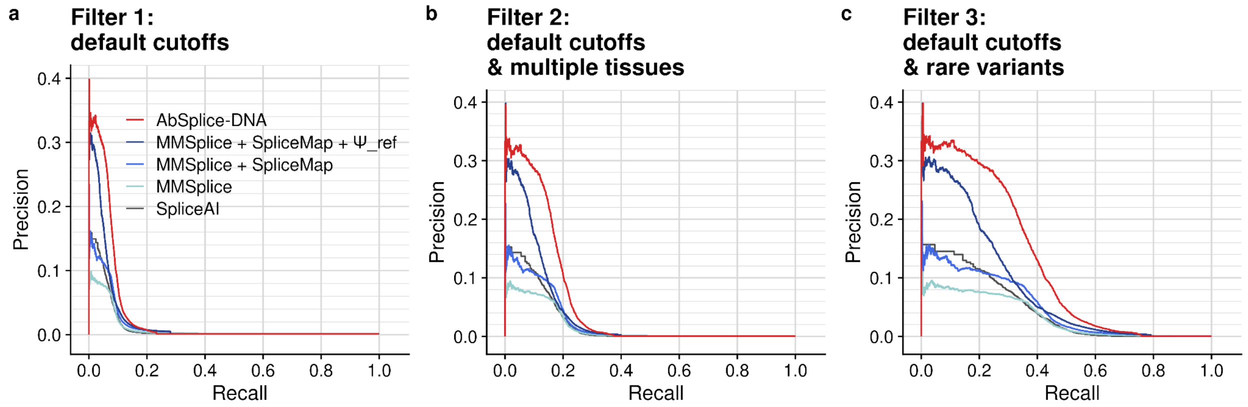


**Supplementary Fig. 2: Performance with different outlier filters.**

Precision-recall curve comparing the overall prediction performance on all GTEx tissues of SpliceAI, MMSplice using GENCODE annotation, MMSplice using SpliceMap annotation, MMSplice using SpliceMap annotation along with quantitative reference levels of splicing, and the integrative model AbSplice-DNA, using different filters for aberrantly spliced genes. **a**, Filter: FRASER default cutoffs ($\Delta\Psi > 0.3$, FDR < 0.05) **b**, Filter: same as (a), but restricting to genes that are aberrantly spliced in at least two different tissues from the same individual. **c**, Filter: same as (a), but restricting to genes that have a rare variant within 100 bp of the splice sites. While the results are best with filter 3, the relative improvements in terms of precision at same recall between the methods is the same as with filter 2. In particular, having restricted to variants 100 nt away from any detected split read boundary (filter 3) did not bias our analysis for the splice-site centric method MMSplice over SpliceAI.

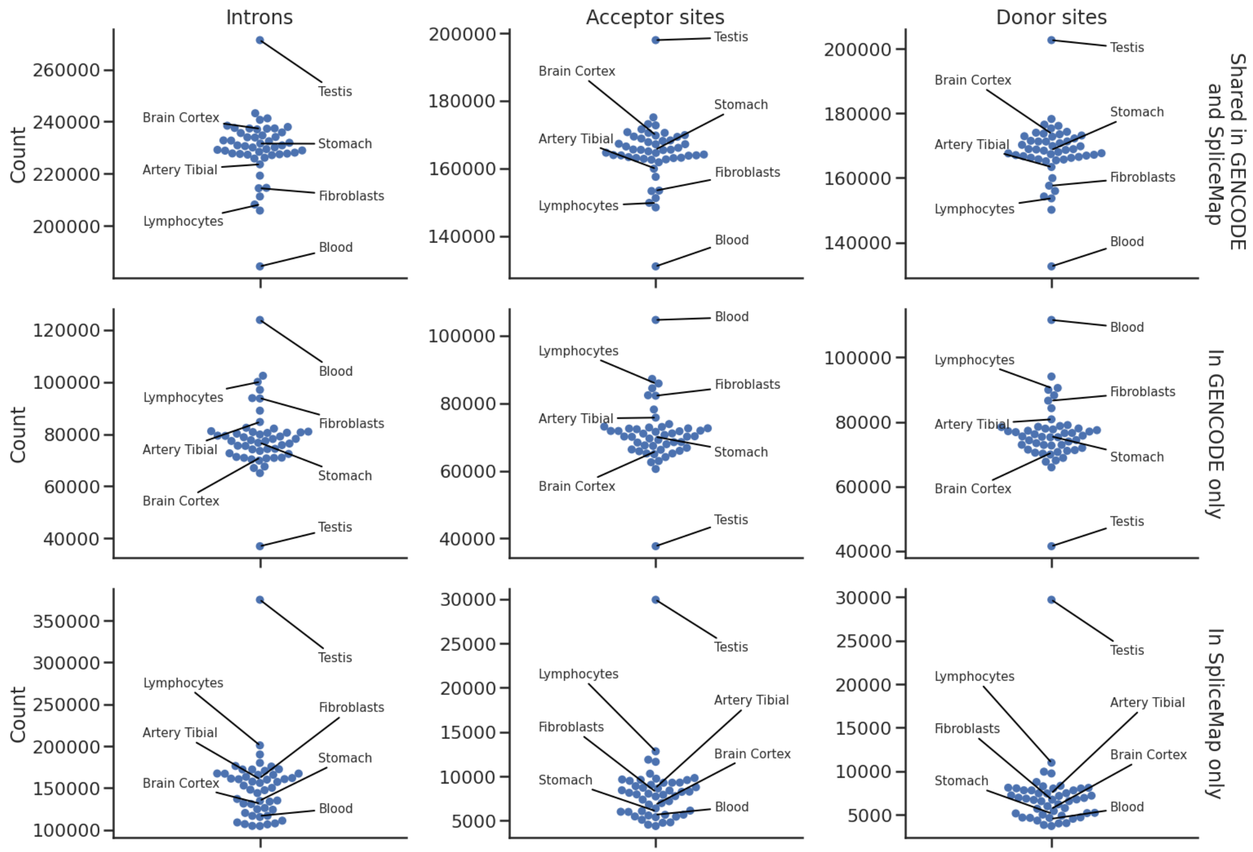


**Supplementary Fig. 3: Comparison of annotated splice-sites in SpliceMap and GENCODE.**

Number of introns, acceptor sites, and donor sites annotated in GENCODE and the SpliceMap of each GTEx tissue (first row), GENCODE only (second row) and SpliceMap only (third row).

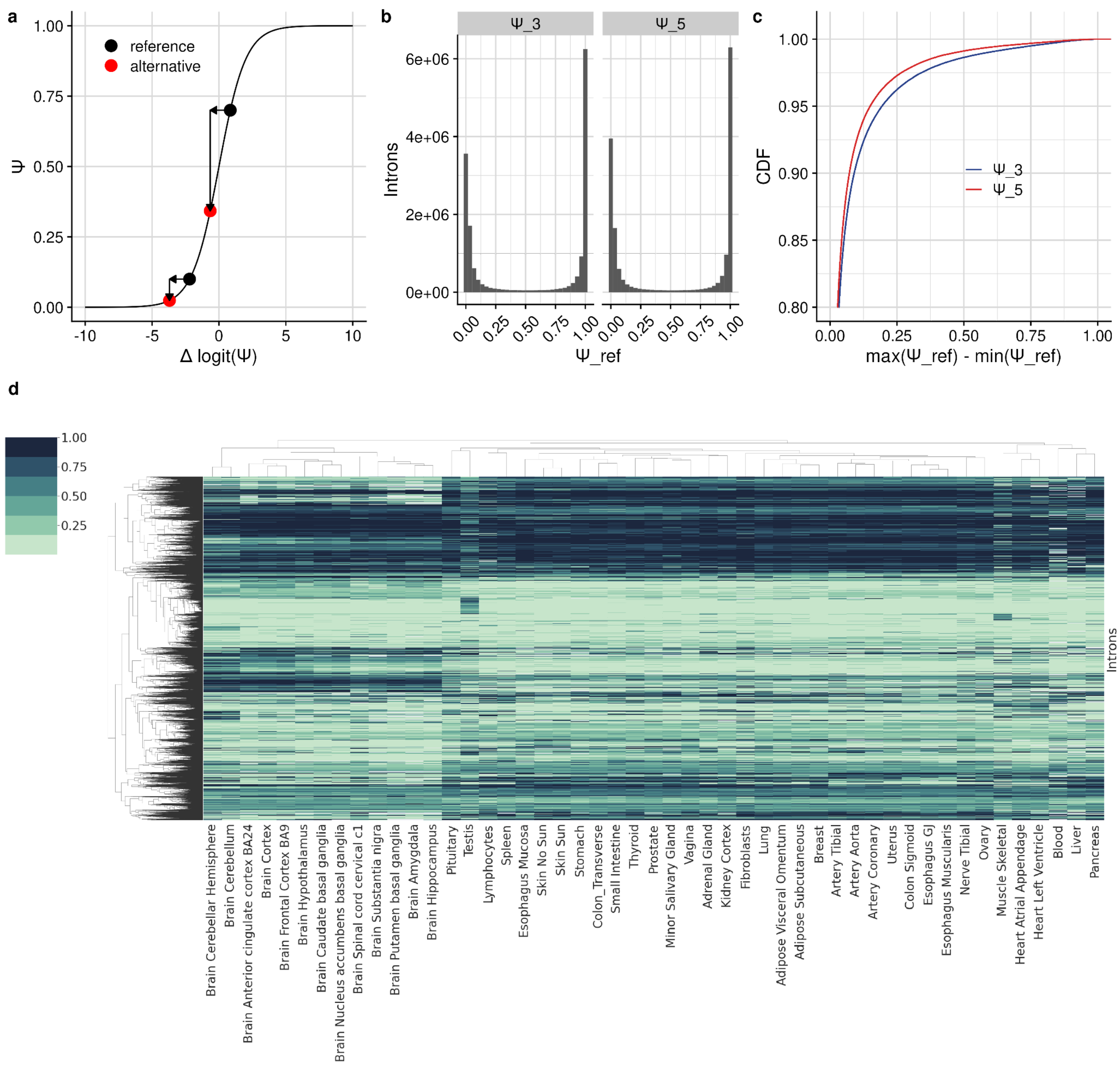


**Supplementary Fig. 4: The variant effect depends on the reference isoform proportion.**
**a**, $\Psi$ against $\Delta logit(\Psi)$ showing the non-linear splicing scaling law. The mutation effect of a variant can lead to different changes in $\Psi$ in natural scale, depending on the reference splicing level of the intron. For example, the same variant can lead to a large change in $\Psi$ if $\Psi_{ref}$ is initially high and almost no change if $\Psi_{ref}$ is initially low. **b**, Distribution of $\Psi_{ref}$ in SpliceMap. Most of the introns are not alternatively spliced, so the reference level of those introns is either 0 or 1. **c**, Cumulative distribution function of the maximum difference of $\Psi_{ref}$ (defined as: $max(\Psi_{ref}) - min(\Psi_{ref})$) across tissues per intron. **d**, Heatmap of the $\Psi_{ref}$ of the most variable introns (defined as: $max(\Psi_{ref}) - min(\Psi_{ref}) > 0.3$) across tissues.

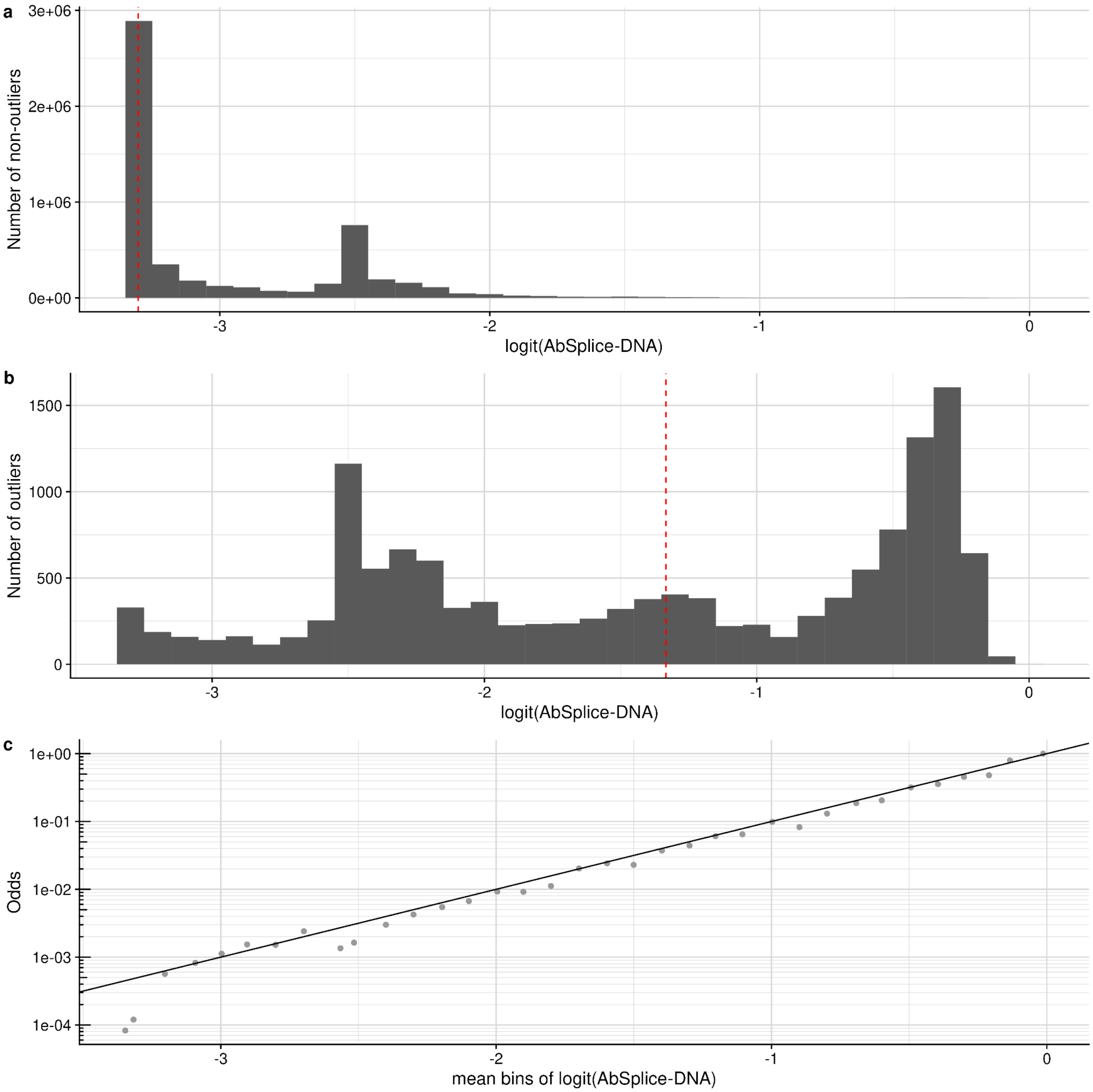


**Supplementary Fig. 5: Calibration of AbSplice-DNA. a**, Histogram of AbSplice-DNA scores for gene, sample, tissue combinations that do not contain an aberrant splicing event. **b**, Histogram of AbSplice-DNA scores for gene, sample, tissue combinations that contain an aberrant splicing event. The peak at logit(AbSplice-DNA) ~-2.5 corresponds to AbSplice-DNA scores that are low due to small SpliceAI and MMSplice scores, but with an expressed splice site as annotated in SpliceMap. The peak at logit(AbSplice-DNA) ~-3.3 corresponds to small SpliceAI and MMSplice scores with an unused splice site as annotated in SpliceMap. **c**, Odds of aberrant splicing events as a function of logit transformed AbSplice-DNA scores (binned in bins of width 0.1). The line represents the diagonal. Note the linear relationship and the (extrapolated) intersection at AbSplice-DNA score of 0.5 (logit(AbSplice-DNA) = 0) corresponding to a log odds of 1, indicating a well calibrated model.

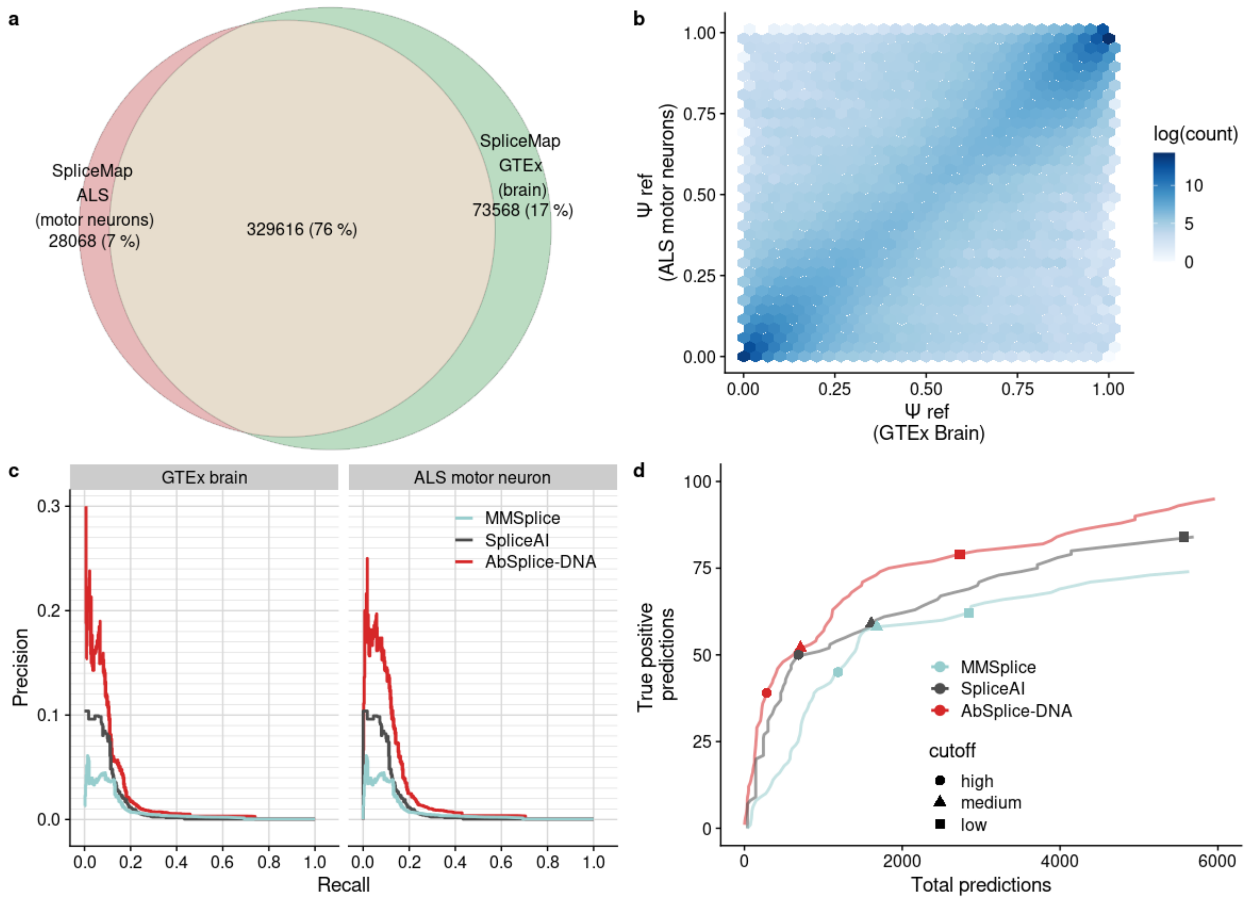


**Supplementary Fig. 6: Replication in the ALS cohort. a,** Venn diagram comparing the total splice sites in the SpliceMaps generated from all different GTEx brain tissues (N=189 ± 39) and the ALS motor neurons (N=45, unaffected individuals of cohort). **b,** Relation of $\Psi_{ref}$ values from GTEx brain SpliceMaps and ALS motor neuron SpliceMap. **c,** Precision-recall curves comparing the prediction performance in the ALS dataset for SpliceAI, MMSplice and AbSplice-DNA with SpliceMaps from GTEx brain tissues and motor neurons. **d,** Predictions causing splicing outliers among total predictions in the ALS dataset at different cutoffs for SpliceAI (low: 0.2, medium: 0.5, high: 0.8), MMSplice (low: 1, medium: 1.5, high: 2), and AbSplice-DNA trained on GTEx and using GTEx brain SpliceMaps (low: 0.01, medium: 0.05, high: 0.2).

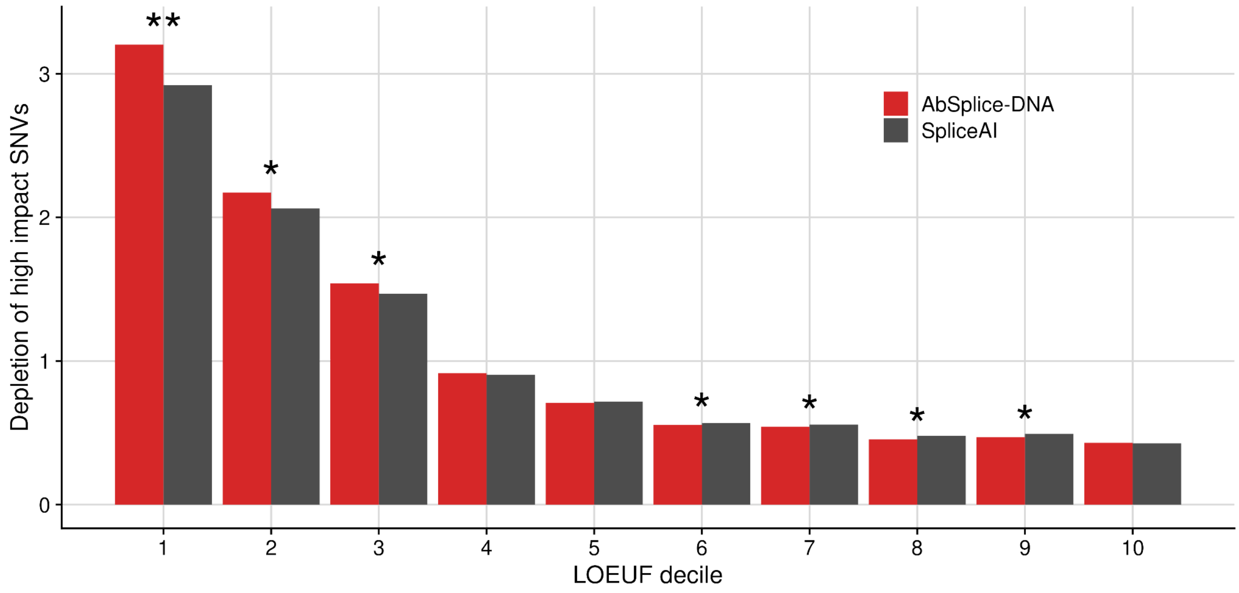


**Supplementary Fig. 7: Depletion of high impact variants in loss-of-function intolerant genes.** Genome-wide depletion of high impact variants among rare SNVs (gnomAD MAF < 0.001) within a gene (N=19,521) as a function of LOEUF score deciles. High impact variants are defined by a SpliceAI cutoff > 0.8 and an AbSplice-DNA cutoff > 0.16, matching the amount of predicted high impact variants of both models (N=109,134). Stars mark significance levels of two-sided Fisher tests (* < 0.05, ** < 10-4).

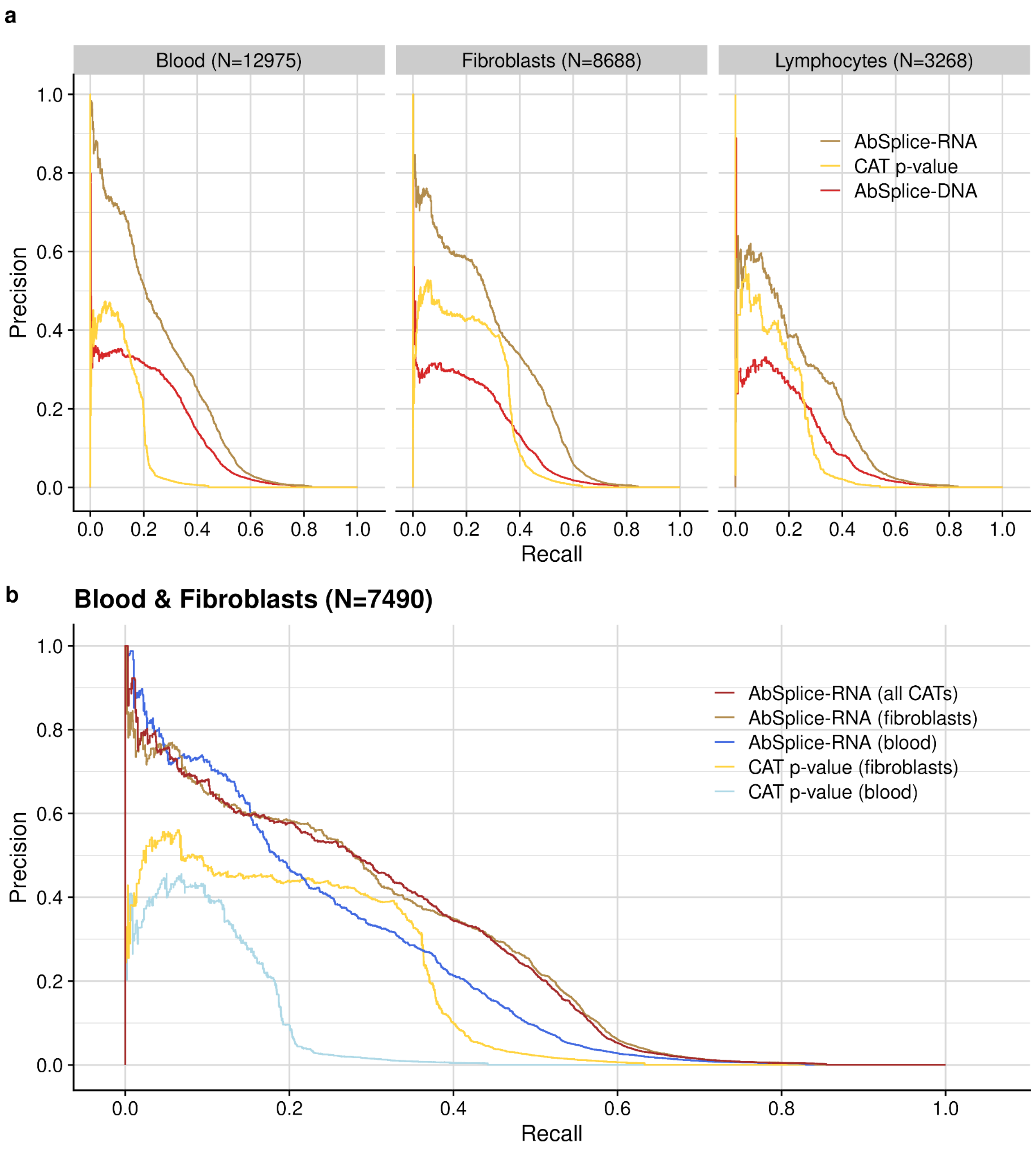


**Supplementary Fig. 8: RNA-based predictions from CAT improve DNA-based scores.**
**a**, Precision-recall curves comparing the overall prediction performance on non-accessible GTEx tissues using the gene-level FRASER p-values from the CAT, AbSplice-RNA trained on a single CAT and AbSplice-DNA. Each panel shows a different CAT and the number of matching samples in the non-accessible tissues. **b**, Same as (a), but for samples having RNA-seq from both blood and fibroblasts. AbSplice-RNA (all CATs) was trained using RNA-seq data from blood, fibroblasts and lymphocytes. Note that AbSplice-RNA (fibroblasts) gave a similar performance as AbSplice-RNA (all CATs). We did not restrict the samples to the ones also having lymphocytes as this would result in a low number of samples (N=2258).
